# T_reg_ cells drive MYCN-mediated immunosuppression and tumor aggressiveness in high-risk neuroblastoma

**DOI:** 10.1101/2022.10.03.510561

**Authors:** Xiaodan Qin, Andrew Lam, Xu Zhang, Satyaki Sengupta, J. Bryan Iorgulescu, Tao Zuo, Alexander E. Floru, Grace Meara, Liang Lin, Kenneth Lloyd, Lauren Kwok, Kaylee Smith, Raghavendar T. Nagaraju, Rob Meijers, Craig Ceol, Ching-Ti Liu, Sanda Alexandrescu, Catherine J. Wu, Derin B. Keskin, Rani E. George, Hui Feng

## Abstract

Solid tumors, especially those with aberrant MYCN activation, harbor an immunosuppressive microenvironment to fuel malignant growth and trigger treatment resistance^1,2^, yet the underlying mechanisms are elusive and effective strategies to tackle this challenge are lacking. Here we demonstrated the crucial role of T regulatory (T_reg_) cells in MYCN-mediated immune repression and tumor aggression using high-risk neuroblastoma (NB) as a model system. Human MYCN-activated NB attracts CD4^+^ T_reg_ cells, which are also found enriched in MYCN-high primary patient samples. Zebrafish *MYCN*-overexpressing neural crests recruit Cd4^+^ cells before tumor formation and induce an immunosuppressive microenvironment, thereby promoting tumor onset and progression. Strikingly, disruption of T_reg_ cells through depletion of forkhead box protein 3a restores anti-tumor immunity and impairs NB development. Together, our studies establish T_reg_ cells as the key driver of MYCN-mediated immunosuppression and tumor aggressiveness, providing mechanistic insights and therapeutic implications.

## Main

Increased activity of the MYCN oncoprotein, through gene amplification, transcriptional upregulation, or protein stability, plays a pivotal role in the initiation, progression, and treatment resistance of various human cancers^3,4^. Besides its intrinsic role in stimulating tumor growth, MYCN also exerts extrinsic effects on the tumor microenvironment (TME) to restrict anti-tumor responses and bolster cancer progression^5,6^. However, the crucial drivers that induce an immunosuppressive TME in MYCN-driven cancers remain unclear. Here we utilize a zebrafish model of MYCN-driven neuroblastoma (NB) together with analyses of patient samples to uncover new mechanisms of MYCN-mediated immunosuppression.

### The zebrafish model of MYCN-driven NB resembles human high-risk disease and harbors an immunosuppressive TME

To understand how MYCN mediates immunosuppression, we studied a zebrafish model of MYCN-driven NB, *MYCN;EGFP*, in which both the human *MYCN* and *EGFP* gene are co-expressed under zebrafish dopamine β hydroxylase (*d*β*h*) promoter^7^. As early as seven weeks post-fertilization (wpf), the *MYCN;EGFP* fish started to develop NB in the interrenal gland, the analogous location of the human adrenal medulla^7^. Despite coming from the same parental fish, the *MYCN;EGFP* siblings had variable disease presentation, ranging from localized tumors to metastatic disease (Fig. 1a). Since *MYCN* amplification is associated with NB aggressiveness^8–10^, we next examined MYCN levels in tumors of these fish. Western blotting analysis revealed an approximately four-fold increase of MYCN protein levels in the metastatic NB compared with the localized tumors (Fig. 1b). These results faithfully reproduced what is observed in human disease, in which high *MYCN* expression is associated with advanced disease and poor prognosis^11^.

**Fig. 1.**
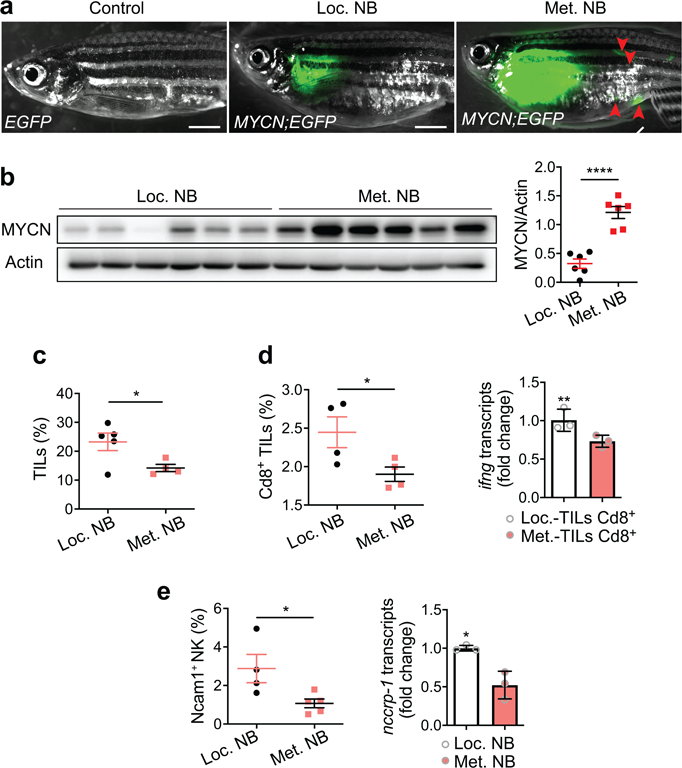
The zebrafish model of MYCN-driven NB resembles human high-risk disease and harbors an immunosuppressive TME. **a**, Overlay of bright field and EGFP images of representative control *Tg*(*d*β*h:EGFP*) and *Tg(d*β*h:MYCN;d*β*h:EGFP)* zebrafish, referred to as *EGFP* and *MYCN;EGFP* respectively, at four months old. EGFP signals enable visualization and quantification of tumors, and red arrowheads point to metastatic tumors. Scale bars, 1 mm. **b**, Western blotting analysis (left) and quantification (right) of MYCN protein levels in neuroblastoma isolated from individual *MYCN;EGFP* sibling fish that developed either localized (Loc.) or metastatic (Met.) tumors at four months of age *(n =* 6). Actin serves as the loading and normalization control. **c-e**, Flow cytometric analysis and quantification of TILs (**c**), Cd8^+^ TILs (**d**), and NK cells (**e**) in localized (Loc.) *vs.* metastatic (Met.) tumors, which expresses low- or high *MYCN*. **d**, **e**, qRT-PCR to detect relative expression of *ifng* in Cd8^+^ TILs (**d**) and the NK marker gene *nccrp-1* in tumors from the above two groups (**e**). *n* indicates the number of biological (**b-e**) or experimental (**d-e**) replicates. Data in (**b-e**) are presented as mean ± SEM and compared for statistical significance by an unpaired two-tailed *t*-test. NS, not significant, *\*P* < 0.05, \*\**P* < 0.01, *\*\*\*\*P <* 0.0001.

In the primary tumor site of metastatic NB patients, *MYCN* amplification is associated with low TILs and inhibits IFN pathways^5,12,13^. To understand whether our zebrafish model also harbors an immunosuppressive TME, we quantified the percentage of TILs in localized and metastatic NB that expressed different levels of MYCN using the published flow cytometric gating strategy^14^. Metastatic NB had a significantly lower percentage of TILs compared to localized NB (Fig. 1c). We next characterized cytotoxic TILs of both groups and found a significantly lower percentage Cd8^+^ and NK cells in *MYCN;EGFP* fish with metastatic tumors compared to those with localized NB (Fig. 1d-e). Consistent with data from human tumor samples^5^, Cd8^+^ TILs from metastatic tumors also showed decreased *ifng* expression (Fig. 1d). The above result is consistent with what was observed in patient *MYCN-*amplified NB^5^, supporting the suitability of the zebrafish model in this research.

### *MYCN*-overexpressing NB attracts Cd4^+^ T_reg_ cells to the TME

The optical transparency of zebrafish enabled us to monitor the dynamic interactions between immune and malignant cells in the living organism during different stages of animal and tumor development^15^. Although *MYCN*-amplified NB is characterized by overall low immune cell infiltration^16^, recent murine studies have revealed the presence of unconventional CD4^+^ cells in the TME^17^. To investigate whether CD4^+^ cells contributed to MYCN-mediated tumor immunosuppression, we bred the *MYCN;EGFP* fish to the *Tg*(*cd4-1:mCherry)* fish to generate *MYCN;EGFP;mCherry* compound transgenic fish, in which neuroblasts and Cd4^+^ cells were labeled with EGFP and mCherry, respectively^18^ (Fig. 2a). Confocal imaging revealed that Cd4^+^ cells were present near the neural crest regions at 1 wpf. By 2 wpf, they began to infiltrate the premalignant mass in *MYCN;EGFP* fish but not the control neural crests of *EGFP* fish (Data not shown). By 3 wpf, a significantly increased number of Cd4^+^ cells surrounded and infiltrated the premalignant mass in *MYCN;EGFP* fish while scarcely encompassing the normal neural crest cells in the control *EGFP* fish (Fig. 2b).

**Fig. 2.**
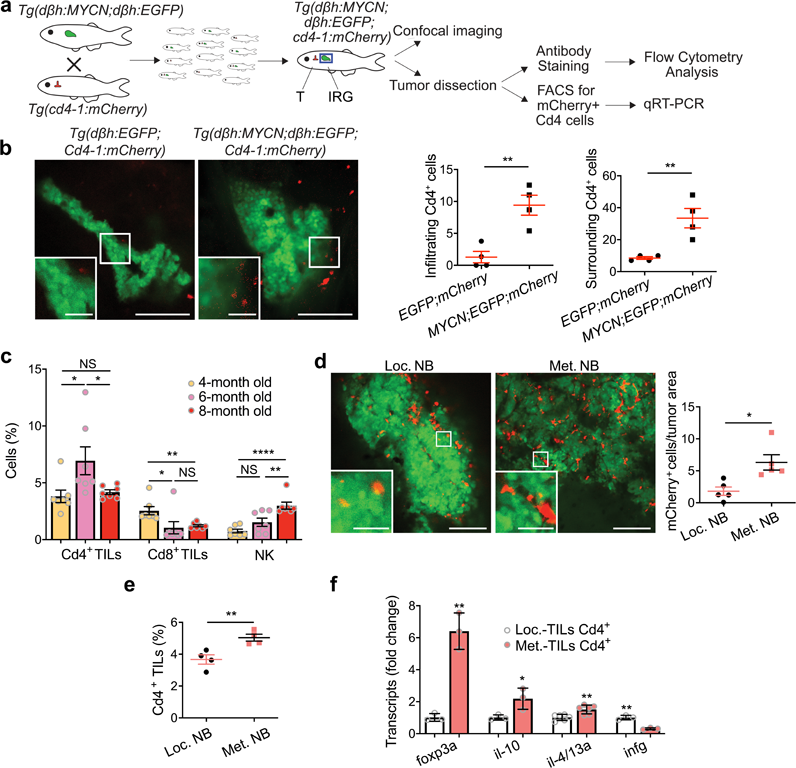
Metastatic NB with high MYCN expression is enriched with immunosuppressive Cd4^+^ T_reg_ cells. **a**, Schematic representation of fish breeding and experimental flow. T: thymus, and IRG: interrenal glands. **b**, Overlay of EGFP and RFP confocal images at the interrenal glands of *Tg(d*β*h:EGFP;Cd4-1:mCherry)* and *Tg(d*β*h:MYCN;d*β*h:EGFP;Cd4-1:mCherry)* fish, referred to as *EGFP;mCherry* and *MYCN;EGFP;mCherry*, respectively. Scale bars = 100 and 25 (inserts) μm. Quantification of Cd4 cells surrounding or infiltrating EGFP non-transformed neural crests or premalignant mass (*n* = 4). **c,** Flow cytometric analysis of Cd4^+^ TIL, Cd8^+^ TIL, and NK cell percentages in the TME of neuroblastoma isolated from fish at the indicated age. **d**, Confocal image overlay showing mCherry^+^ Cd4 cells and EGFP^+^ tumor cells in localized (Loc.) *vs.* metastatic (Met.) tumors of four-month-old zebrafish (left panels). Quantification of mCherry^+^ Cd4 cells per tumor area (center*; n* = 5). **e**, Flow cytometric analysis of Cd4^+^ TILs in localized *vs.* metastatic tumors from fish at 6-month-old (right, *n* = 4), which expresses low- or high *MYCN*, respectively. **f**, qRT-PCR to detect relative expression of *foxp3a*, *il10*, *il-4/13a*, and *ifng* in Cd4^+^ TILs from localized and metastatic tumors (*n =* 3 for all groups except *il-4/13a* [*n =* 6]). *n* indicates the number of biological (**b**-**e**) or experimental (**f**) replicates. Data are representative of two experiments (**b**-**e**). Data are presented as mean ± SEM and compared for statistical significance by an unpaired two-tailed *t*-test. NS, not significant*, *P* < 0.05, \*\**P* < 0.01, *\*\*\*\*P <* 0.0001.

To monitor TIL dynamics in the TME during tumor development, we analyzed *MYCN;EGFP* fish of different ages, with older fish developing more aggressive disease. Cd4^+^ cells comprised a larger proportion of TILs than Cd8^+^ and NK cells (Fig. 2c). Interestingly, the percentages of TILs fluctuated as fish aged and tumors progressed, with Cd4^+^ cells peaking at 6 months and declining at 8 months (Fig. 2c). As expected, frequencies of both Cd8^+^ and NK cells in the TME were low; however, Cd8^+^ cells declined continuously while NK cells rose moderately with age (Fig. 2c). These results indicate a developmental oscillation of TILs during tumorigenesis.

Since Cd4^+^ cells peaked in tumors around 6-month-old fish, we analyzed fish at 4 and 6 months old to determine whether *MYCN* impacts Cd4^+^ infiltration into the TME. Confocal imaging revealed more mCherry+ cells inside the TME of zebrafish with localized NB compared to those with metastatic NB. Quantification of these mCherry^+^ cells showed a significantly more Cd4^+^ cells in metastatic NB (Fig. 2d). Flow cytometric analysis also revealed a striking increase of Cd4^+^ TILs in these metastatic tumors (Fig. 2e). To understand the composition of these Cd4^+^ TILs, we examined a panel of differentiation marker genes. We found that Treg signature genes *foxp3a* and *il-10* were significantly upregulated in Cd4^+^ TILs isolated from metastatic NB compared to localized NB (Fig.2f). In contrast, the transcript levels of *ifng* (Th1 marker gene) and *il-4-13a* (Th2 marker gene) were downregulated (Fig.2f). Taken together, our results demonstrate that the aggressive NB with elevated MYCN expression is enriched with Cd4^+^ Tregs.

### T_reg_ disruption impairs *MYCN*-driven NB development

The transcriptional factor FOXP3 is critical for the development and immunosuppressive functions of T_reg_ cells^19,20^. In zebrafish, homozygous *foxp3a* mutations disrupt the development and function of T_reg_-like cells^21^. Our findings show that Cd4^+^ T_reg_ cells preferentially infiltrate the metastatic NB (Fig. 2). We thus examined the role of T_reg_ cells in NB pathogenesis. Specifically, we bred our transgenic *MYCN;EGFP* fish to *foxp3a* (um252/+) heterozygous fish, which develop and survive similarly to wild-type fish^21^, yet have reduced T_reg_-like cells (Extended Data Fig. 1a). The resulting progeny were subsequently raised and monitored for tumor development as described^7^. Although fish from both groups began to develop NB at 7 wpf, there were fewer *MYCN;EGFP* fish with allelic loss of *foxp3a* developed tumors compared to their wild-type siblings (Fig. 3a-b). By 20 wpf, 73% of *MYCN;EGFP* fish had already developed tumors, while there were only 54% of *MYCN;EGFP;foxp3a+/-* fish with tumors (Fig. 3b).

**Fig. 3.**
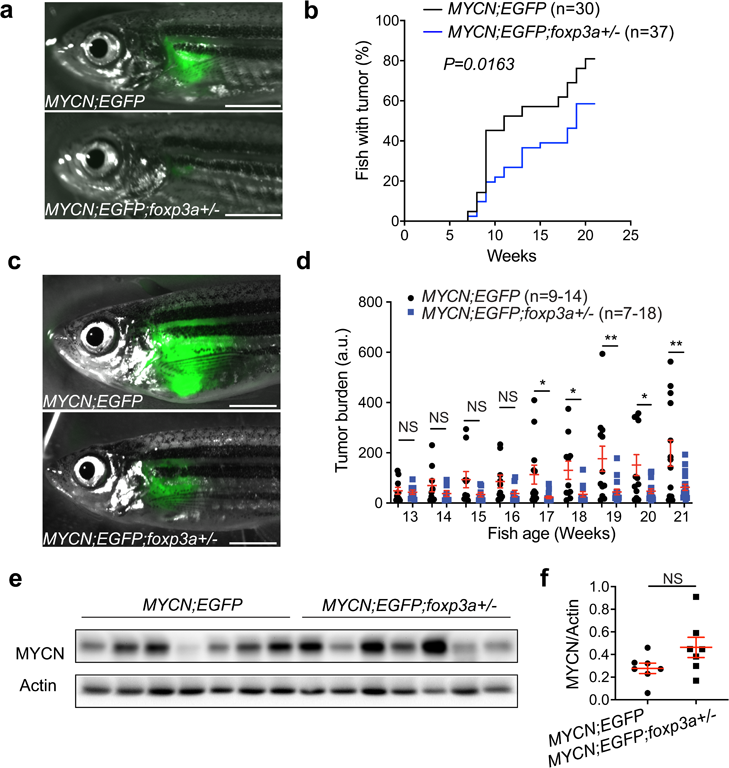
Allelic loss of *foxp3a* in zebrafish decreases MYCN-driven NB aggressiveness. **a**, Overlay of brightfield and EGFP images of a *MYCN;EGFP* (top) and *MYCN;EGFP;foxp3a+/-* (bottom) sibling fish at seven weeks of age. Scale bars = 2 mm. **b**, Kaplan-Meier analysis showing the rate of tumor onset in *MYCN;EGFP* (black line, n=30 per group) and *MYCN;EGFP;foxp3a+/-* (blue line, n=37) sibling fish. Statistical analysis was performed by the log-rank Mantel-Cox test. **c**, Overlay of brightfield and EGFP images of a *MYCN;EGFP* (top) and *MYCN;EGFP;foxp3a+/-* sibling fish (bottom) at 17 weeks of age. Scale bars = 1 mm. **d**, Quantification of tumor burden over time based on EGFP intensity in *MYCN;EGFP* fish and their age-matched *MYCN;EGFP;foxp3a+/-* siblings (n=9-14 and 7-18, respectively). **e**, Western blotting analysis (**e**, left) and quantification (right) of MYCN protein levels in EGFP^+^ neuroblastoma cells isolated from the individual *MYCN;EGFP* and *MYCN;EGFP;foxp3a+/-* fish at six-month-old (*n =* 7). *n* indicates the number of biological replicates (**b**, **d**, **e**). Data are presented as mean ± SEM and compared to an unpaired two-tailed *t*-test. NS, not significant, *\*P* < 0.05, \*\**P* < 0.01.

Next, we asked whether the allelic loss of *foxp3a* could impact NB progression. Fluorescent microscopy imaging of the two groups showed similar tumor burden during the first few weeks of tumor onset; however, tumors in *MYCN;EGFP* fish grew faster than those in *MYCN;EGFP;foxp3a+/-* fish (Fig. 3c-d). By four weeks post tumor onset, *MYCN;EGFP* fish had a significantly more tumor burden than their *MYCN;EGFP;foxp3a+/-* siblings (Fig. 3d). Importantly, *foxp3a* loss did not impact *MYCN* expression in NB in the two fish groups, as demonstrated by similar MYCN protein levels in tumor cells (Fig. 3e). Collectively, our data demonstrate that T_reg_ cells play a critical role in promoting NB initiation and progression.

### T_reg_ disruption restores anti-tumor immune responses in MYCN-driven NB

To understand the mechanism by which T_reg_ promotes NB pathogenesis, we next investigated the effect of disrupting T_reg_ through allelic loss of *foxp3a* on tumor immunosurveillance. Tumors from *MYCN;EGFP* and *MYCN;EGFP;foxp3a+/-* fish were dissected and dissociated. The percentage of total TILs, Cd4^+^, Cd8^+^ T cells, and NK cells were then quantified by flow cytometric analysis. Consistent with their less aggressive tumor phenotypes (Fig. 3c-d), *MYCN;EGFP;foxp3a+/-* fish had fewer EGFP^+^ tumor cells compared to *MYCN;EGFP* fish (Extended Data Fig. 1b). Importantly, disrupting T_reg_ cells in *MYCN;EGFP;foxp3a+/-* fish led to more TILs than *MYCN;EGFP* fish (Fig. 4a). In addition, significantly higher frequencies of Cd4^+^ and Cd8^+^ TILs but not NK cells were detected in *MYCN;EGFP;foxp3a+/-* fish than *MYCN;EGFP* fish (Fig. 4b-e). Cd4^+^ TILs from *MYCN;EGFP;foxp3a+/-* tumors were more immunoreactive than those from *MYCN;EGFP* tumors, as demonstrated by reduced *il-10* and increased *ifng* expression (Fig. 4d). Moreover, Cd8^+^ TILs from *MYCN;EGFP;foxp3a+/-* tumors showed upregulated *ifng* expression, indicating improved cytotoxicity (Fig. 4e). Interestingly and in contrast to the TME, the percentage of Cd4^+^ cells was similar in the kidneys among these two groups of fish, indicating that the enhancement of anti-tumor immune responses is tumor-specific (Extended Data Fig. 1c). Taken together, these findings indicate that anti-tumor immune responses are restored in MYCN-driven NB when T_reg_ development and functions are disrupted through genetically depleting Foxp3a^21^.

**Fig. 4.**
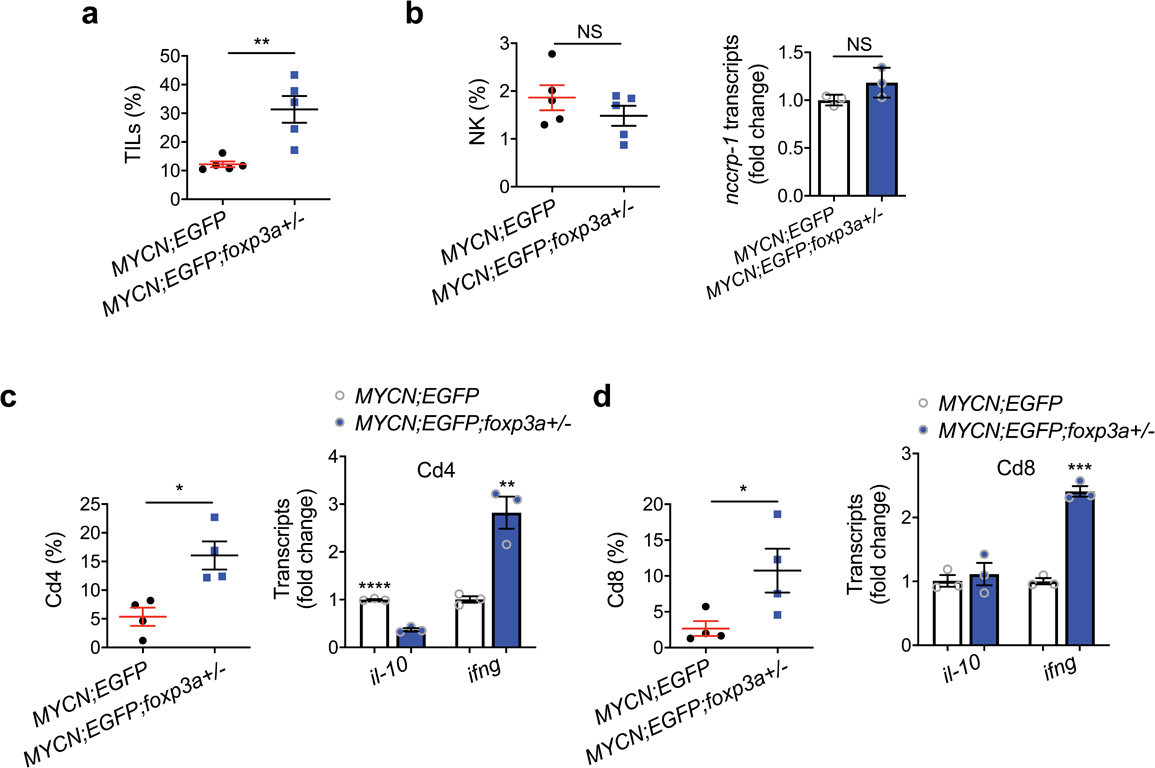
Treg disruption restores anti-tumor immune responses in MYCN-driven NB. **a**, Flow cytometric analysis and quantification of TILs (a), NK cells (**b**), Cd4^+^ TILs (**c**), and Cd8^+^ TILs (**d**) in the TME of NB from *MYCN;foxp3a+/+* and *MYCN;foxp3a+/-* sibling fish at six months of age (*n =* 5). **b**-**d,** qRT-PCR analysis of the NK marker gene *nccrp-1* (**b**), as well as *il10* and *ifng* from Cd4^+^ (**c**, left) and Cd8^+^ (**d**, right) TILs, from the two fish groups (*n =* 3). *n* indicates the number of biological (**a-d**) or experimental (**b**-**d**) replicates. Data are presented as mean ± SEM and analyzed for statistical significance by an unpaired two-tailed *t*-test. NS, not significant; * *P* < 0.05; ** *P* < 0.01; **** *P* < 0.0001.

### Human MYCN-activated NB attracts Tregs

To understand whether human MYCN-activated NB cells can attract CD4^+^ T_reg_ cells to the TME, we applied an *in vitro* system to monitor the interactions between human NB and T cells (Fig. 5a). We utilized human SHEP-MYC-ER NB cells, in which 4-hydroxytamoxifen (4-OHT) treatment conditionally regulate MYCN activation^22^. In co-culture assays, conditioned media from 4OHT-treated SHEP-MYCN-ER cells led to significantly more recruitment of CD4^+^ and T_reg_ but not CD8^+^ T cells (Fig. 5b).

**Fig. 5.**
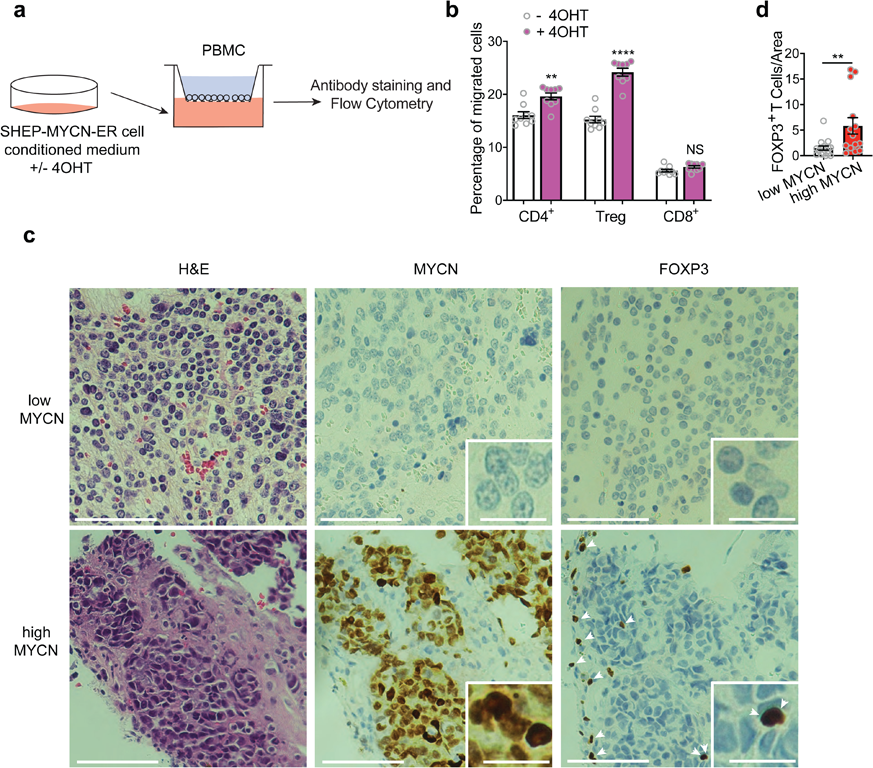
Human MYCN-activated NB attracts CD4^+^ FOXP3^+^ T_reg_ cells to the TME. **a**, Schematic drawing of the transwell migration assay: media from SHEP-MYCN-ER cells treated with or without 4OHT were added into the lower chamber of a transwell migration plate, with human PBMCs in the top chamber. **b,** Percentages of human CD4^+^, T_reg_, and CD8^+^ T cells that migrated to the lower chamber of a transwell plate, containing media of human SHEP-MYCN-ER cells treated with or without 4OHT for 48 hours (*n* = 9). **c**, Representative histological and immunohistochemical staining (brown) of MYCN (nuclear) and FOXP3 (nuclear) in clinical neuroblastoma patient samples. Inserts are zoom-in views of tumor cells together with or without FOXP3^+^ T cells. Arrowheads point to FOXP3^+^ T cells. Scale bars = 200 and 50 (inserts) μm. **d**, Quantification of FOXP3^+^ cells per tumor area (*n* = 16 and 14 for MYCN low and high tumor samples).

To assess the patient relevance of these findings, we performed immunohistochemical staining to detect MYCN and FOXP3 protein expression in primary NB samples. As expected, MYCN protein levels were significantly more in tumors with *MYCN* amplification (Extended Data Fig. 2). Consistent with our findings in the zebrafish model, FOXP3^+^ T cells were significantly enriched in tumors with high MYCN expression compared to low MYCN-expressing samples, indicating the recruitment of T_reg_ cells by MYCN (Fig. 5c-d, Extended Data Fig. 3, Supplementary Table 2). We also observed FOXP3^+^ T cells surrounding the tumor mass, which likely form a barrier to prevent the entry of cytotoxic CD8^+^ and NK cells into the TME (Fig. 5c). Taken together, human MYCN-activated NB cells recruit FOXP3^+^ T_reg_ cells to mediate immunosuppression and tumor aggression.

Immunologically “cold” tumors, such as those MYCN-driven NB, are challenging to treat and are invariably associated with poor prognosis^23–25^. Despite the knowledge that *MYCN* amplification is associated with low frequencies of tumor-infiltrating B cells, T cells, NK cells, and M1 macrophages^16,26,27^, how MYCN induces a broadly immunosuppressive TME remains unknown. Here, we demonstrate that zebrafish metastatic NB with high-MYCN expression behaves similarly to human high-risk NB, displaying fewer TILs, especially cytotoxic Cd8^+^ and NK cells. Our studies highlight the conservation of the zebrafish in both oncogenic signaling and immune system development, demonstrating this organism to be both a suitable and advantageous *in vivo* system for tumor immunology research. Importantly, an increased frequency of T_reg_ cells was found to infiltrate metastatic NB with high-MYCN expression in zebrafish, suggesting that the upregulation of MYCN is critical for T_reg_ recruitment to the TME and their immunosuppressive functions^28,29^. Moreover, the ability of MYCN to promote high-risk NB is dependent on T_reg_ cells, as disrupting T_reg_ through *foxp3a* loss significantly delays NB onset and reduces tumor burdens. Hence, our studies identify the MYCN-Treg axis as an inducer of the immunosuppressive TME in high-risk NB.

Although the presence of T_reg_ cells in the TME is a characteristic feature of human tumors, its role in cancer is context-dependent. For example, increased infiltration of T_reg_ cells into the TME is associated with unfavorable outcomes in melanoma, cervical cancer, and renal cancer^30–32^, while predicting better patient survival in colorectal cancer and head and neck cancer^33,34^. Although T_reg_ cells were found in the bone marrow of patients with *MYCN*-amplified NB and a chimeric MYCN/ALK^F1174L^ mouse model of NB^35,36^, it was unclear whether they contribute to immunosuppression and tumor aggression in NB. By using the zebrafish, with a developing immune system orchestrating with tumorigenesis as in NB children, we show that T_reg_ cells preferentially infiltrate the TME of metastatic NB with high-MYCN expression. Importantly, through the use of a *foxp3a* heterozygous fish, our study establishes the functional importance of T_reg_ cells in promoting MYCN-driven NB onset and progression through the induction of an immunosuppressive TME. Hence, besides boosting tumor cell growth and proliferation, MYCN can also foster NB development by inducing T_reg_-mediated immunosuppression. T_reg_ cells have previously been shown to exert a broad negative effect on TILs, macrophages, and neutrophils^16,27^. Our findings that high-MYCN NB harbors increased T_reg_ cells explain the clinical observation that *MYCN-*amplified NB has an unusually low number of immune cells of various types^16^.

Due to poor baseline immune responses in the TME of NB, ∼30% of children relapse during or after treatment with current immunotherapy (e.g., anti-GD2 monoclonal antibodies)^37^. T_reg_ cells have been proposed as an immunological target for multiple cancer types. Depletion of tumor-resident Foxp3^+^ T_reg_s has been shown to elicit effective anti-tumor immunity in lung adenocarcinoma, lymphoma, and pancreatic cancer^38–40^. Although clinical approaches targeting T_reg_ cells show some efficacy, it is challenging to 1) specifically target tumor-resident T_reg_ cells without causing severe autoimmunity, and 2) deplete T_reg_ cells without impacting other tumor-infiltrating effector cells. Identifying and disturbing the molecular connection between MYCN and T_reg_ cells could serve as an alternative approach to target tumor-resident T_reg_ cells without causing high toxicities to treat MYCN-driven high-risk NB and perhaps other MYCN-driven cancers as well.

## Methods

### Zebrafish husbandry

Zebrafish husbandry and handling were performed in the aquatic facility at Boston University School of Medicine under approved protocols from the Institutional Animal Care and Use Committee (IACUC). The *Tg(d*β*h:EGFP)* and *Tg(d*β*h:MYCN;d*β*h:EGFP)* fish in the AB background were obtained from the A. Thomas Look Laboratory at Dana-Farber Cancer Institute^41^. The *Tg(foxp3a:EGFP)* and *foxp3a+/-* fish in the AB background were obtained from Dr. Craig Ceol’s laboratory at the University of Massachusetts Worcester^21^. The *Tg(Cd4-1:mCherry)* fish in the AB background were obtained from Dr. Adam Hurlstone’s laboratory at the University of Manchester^18^.

### Zebrafish genotyping and tumor surveillance

Progenies were raised and screened at 5 days post-fertilization (dpf) under a fluorescent dissecting microscope (Olympus) for EGFP-expressing cell masses at the interrenal gland (IRG) to confirm the presence of *MYCN* transgene. At 4 wpf, the sorted fish were fin-clipped to extract genomic DNA for PCR amplifications using gene-specific primers and Taq DNA polymerase (New England Biolabs). The following primers were used to identify fish harboring *foxp3a* mutations: forward 5’-CAGTTCTGAAGGCAAAGGG-3’ and reverse 5’-AATCGGTGCTTATGCGTC-3’ as previously described^21^. Fish were monitored and imaged weekly for tumor development as described^7,41^. Tumor burden was quantified using ImageJ software (National Institute of Health, NIH).

### Frozen sectioning and confocal microscopy

Fish were euthanized as described in IACUC protocols, subsequently fixed in 4% paraformaldehyde (ThermoFisher) at 4°C overnight with gentle agitation, washed with phosphate-buffered saline containing 0.1% Tween 20 (Fisher), equilibrated in 30% sucrose at 4°C overnight, and frozen at −80°C. The frozen fish were embedded in O.C.T compound (SAKURA) and sectioned at a 14 µm thickness using a cryostat (ThermoFisher).

Live embryos and fish embedded in 1% low-melting agarose (Life Technologies) or fixed samples were imaged using an LSM 710-Live Duo Confocal microscope (Zeiss). Images were processed using ImageJ software (NIH).

### Flow cytometry

Tumors were dissected under a fluorescent microscope (Olympus), dissociated in a mixture of RPMI 1640 (Corning), 0.025 μg/ml Liberase (Roche), 0.6μg/ml DNase I (ThermoFisher), and 1x Pen/strep (Corning), washed in RPMI 1640 supplemented with 10% fetal bovine serum (FBS; Sigma), and filtered with a 40-μm filter (Falcon). Fish kidneys were dissected, dissociated, and filtered in RPMI 1640 supplemented with 10% FBS. Single-cell suspensions were stained with the primary antibodies: rat anti-fish CD4 (1:100, Clone 6D1, Bio Cosmo, CAC-NIH-NA-01), CD8 (1:100, Clone 2C3, Bio Cosmo, CAC-NIH-NA-02)^42–44^, or anti-zebrafish Ncam1 antibody (1:100, Institute for Protein Innovation) in PBS containing 10 U/mL Heparin (Sigma), 10% FBS, and 1x Pen/strep, for 30 min at 4°C. Cells were then stained with the secondary antibody, goat anti-rat APC (1:500; Invitrogen, A10540) or goat anti-human (1:200; Jackson Immuno, 109-136-170) for 30 min at 4°C, and counterstained with DAPI (1:5,000, ThermoFisher, 62248). Analysis was performed on an LSRFortessa flow cytometer (BD Biosciences), and Fluorescent-Activated Cell Sorting (FACS) was performed on a FACSAria II (BD Biosciences). Purified cells were used for RNA extraction and qRT-PCR analysis. FlowJo software 10.4 (Tree Star) was used to analyze all flow cytometric data.

### Quantitative real-time-PCR (qRT-PCR)

RNA was extracted from cells using Trizol (Fisher) and/or the Rnaeasy microKit (Qiagen), and subjected to gDNA clean-up and cDNA synthesis with QuantiTect Reverse Transcription Kit (Qiagen). SYBR Green PCR master mix (Applied Biosystems) and a Step-One PCR instrument (Applied Biosystems) were utilized for the qRT-PCR reaction. The qRT-PCR primers are listed in **Supplementary Table 2**. *ACTB* or *act* expression levels were used to normalize the differences in RNA input. All reactions were performed in triplicates.

### Western blotting

Tumors were lysed in RIPA buffer (1% NP-40, 0.1% SDS, 50 mM Tris-HCl pH 7.4, 150 mM NaCl, 0.5% sodium deoxycholate, and 1 mM EDTA) supplemented with 1x or 2x Halt protease and phosphatase inhibitor cocktail (ThermoFisher). Primary antibodies included anti-MYCN (1:1000; Santa Cruz Biotechnology, sc-53993) and anti-Actin (1:1000; Santa Cruz Biotechnology, sc-47778). Secondary antibodies included horseradish-peroxidase-conjugated anti-mouse or anti-rabbit IgG (1:2000 or 1:5000; ThermoFisher, 31430 or 31460). Proteins were detected by ECL chemiluminescence detection kits (ThermoFisher) and autoradiographs were obtained with a G:BOX Chemi XT4 (Syngene) and a CCD camera. Quantification analysis was performed using Syngene GeneTools software (Syngene).

### Transwell migration assay

Media from human SHEP-MYCN-ER cells treated with or without 4OHT for 48 hours were placed in lower chambers of the transwell plates with a 5-μm pore membrane (Corning). Peripheral blood monocytes (PBMCs) were isolated from human blood (Research Blood Components LLC.) through centrifugation with the Ficoll Pague reagent (Cytiva) at 400g for 30 min, with acceleration/deceleration rates at 9/0. PBMCs were briefly incubated in RPMI 1640 media supplemented with 10% FBS for 2 hours to collect nonadherent cells, which were added to the upper chamber of the transwell plates at a density of 10^6^ cells/100 µl. The plates were incubated at 37°C supplemented with 5% CO_2_. After 4 hours, cells from both upper and lower chambers were collected and stained with BV421-conjugated CD4 (1:100, Clone RPA-TA, BioLegend, 300501) and APC/CY7-conjugated CD8 (1:100, Clone SK1, BioLegend, 344702), as well as PE-conjugated anti-CD25 (1:100, Clone M-A251, Biolegend, 356101). These cells were subsequently fixed and permeabilized with a Human FOXP3 buffer set (BD Biosciences) and then stained with Alexa Fluor 647 conjugated anti-FOXP3 mAbs (1:100, Clone 259D/C7, BD Biosciences, 560044) to identify CD4^+^CD25^+^FOXP3^+^ T_reg_ cells. The percentages of migrated cells were calculated by dividing the number of migrated cells in the lower chamber by the total cells of the same type.

### Immunohistochemical staining and scoring

Formalin-fixed, paraffin-embedded tumor tissue was obtained from neuroblastoma patients that had targeted multigene sequencing at Brigham and Women’s Hospital/Dana Farber Cancer Institute^45^. Patient samples were analyzed under Mass General Brigham Institutional Review Board approval (#2019P000017). 4 µm-thick tissue sections were used for hematoxylin and eosin staining and immunohistochemical staining, the latter of which was performed using the Leica Bond III automated staining platform and Leica Biosystems Refine Detection Kit. The primary antibodies include anti-N-MYC (1:160 dilution, clone D4B2Y, Cell Signaling Technology, 51705; citrate antigen retrieval) and anti-FOXP3 (1:50 dilution, clone D2WE8, Cell Signaling Technology, 98377). The staining was analyzed by board-certified pathologists who were blinded to the samples’ molecular status. Nuclear MYCN staining was assessed under 200X magnification, using a modified H-score (ranging from 0-300), in which the percent of tumor cells with positive staining was multiplied by the staining intensity (graded from 0 none to 3 strong)^46^. The number of FOXP3^+^ T cells was quantified at 200x magnification, divided by the field of view area to obtain cell density, and then averaged across multiple fields of view.

### Statistical analysis

Statistical analysis was performed with GraphPad Prism 8.0 using unpaired two-tailed t-tests unless otherwise stated. Patient survival and zebrafish tumor onset curves were estimated using Kaplan-Meier methods and compared using the log-rank Mantel-Cox test. NS, not significant *P* >= 0.05, *\*P* < 0.05, \*\**P* < 0.01, *\*\*\*P <* 0.001, *\*\*\*\*P <* 0.0001.

## Data availability

All raw data are available upon request by emailing the corresponding author Dr. Hui Feng at

## Material availability

The zebrafish NCAM1 antibody—IPI-zNCAM1—used as a marker for NK cells, is available for research studies from The Institute for Protein Innovation. For more information, contact.

## Acknowledgments

We thank Drs. Chuan Yan, David M. Langenau, Nicole M. Anderson, and M. Celeste Simon for reagents; Kelly Miao for technical assistance; Drs. Yi Zhou and Leonard Zon for helpful discussions; Dr. Shuaiying Cui for equipment; Dr. Michael T. Kirber from the Imaging Core Facility at Boston University School of Medicine for his expert help with imaging acquisitions; Dr. Ronald Mathieu from Boston Children’s Hospital Flow Cytometry Core Facility for assistance with cell sorting, and Caitlin Edwards and the Dana-Farber/Harvard Cancer Center Specialized Histopathology Core for immunohistochemistry services. This study was supported by grants from the National Institutes of Health (NIH: CA134743 and CA215059), the St. Baldrick’s Foundation, the National Science Foundation (1911253), the American Cancer Society (RSG-17-204-01-TBG), and the Alex’s Lemonade Stand Foundation to H.F; X.Z. is supported by a postdoctoral fellowship from the China Scholarship Council (No. 201808440648).

S.S. is supported by a Young Investigator Award from the Rally Foundation for Childhood Cancer Research and the Friends for Life Neuroblastoma Fellowship. J.B.I. acknowledges support from the NIH (K12CA090354) and Conquer Cancer Foundation. A.F. and K.S. are supported by the Undergraduate Research Opportunity Program from Boston University. D.B.K. is supported by a grant from the NIH (R01HL157174). R.G. acknowledges support from the St Baldrick’s Foundation, a DoD Idea Award (CA191000), and an NIH grant (R01CA271605). The Dana-Farber/Harvard Cancer Center is supported in part by an NCI Cancer Center Support Grant (NIH P30CA06516). The content of this research is solely the responsibility of the authors and does not necessarily represent the official views of the NIH.

## Author contributions

H.F. and X.Q. conceived the project. H.F., X.Q., A.L., X.Z., S.S., and R.E.G. designed the experiments. X.Q., A.L., X.Z., G.M., A.F., K.S., S.S., and L.K. performed the experiments. X.Q., X.Z., A.L., S.D., A.F., S.S., L.K., T.Z., and H.F. analyzed the data. J.B.I., T.Z., and S.A. designed, implemented, and analyzed the immunohistochemistry experiments. K.L., R.M., C.C., D.B.K., C.W., and R.T.N. provided reagents and advice. X.Q., H.F., X.Z., G.M., A.F., L.K., D.V., and R.E.G. wrote the manuscript with inputs from all other authors. H.F. and R.E.G. supervised the project. R.E.G., S.S., C.C., D.B.K., J.B.I., L.L., C.T.L., C.W., and R.T.N. provided experimental support and intellectual inputs.

---

**Extended Data Fig. 1.**
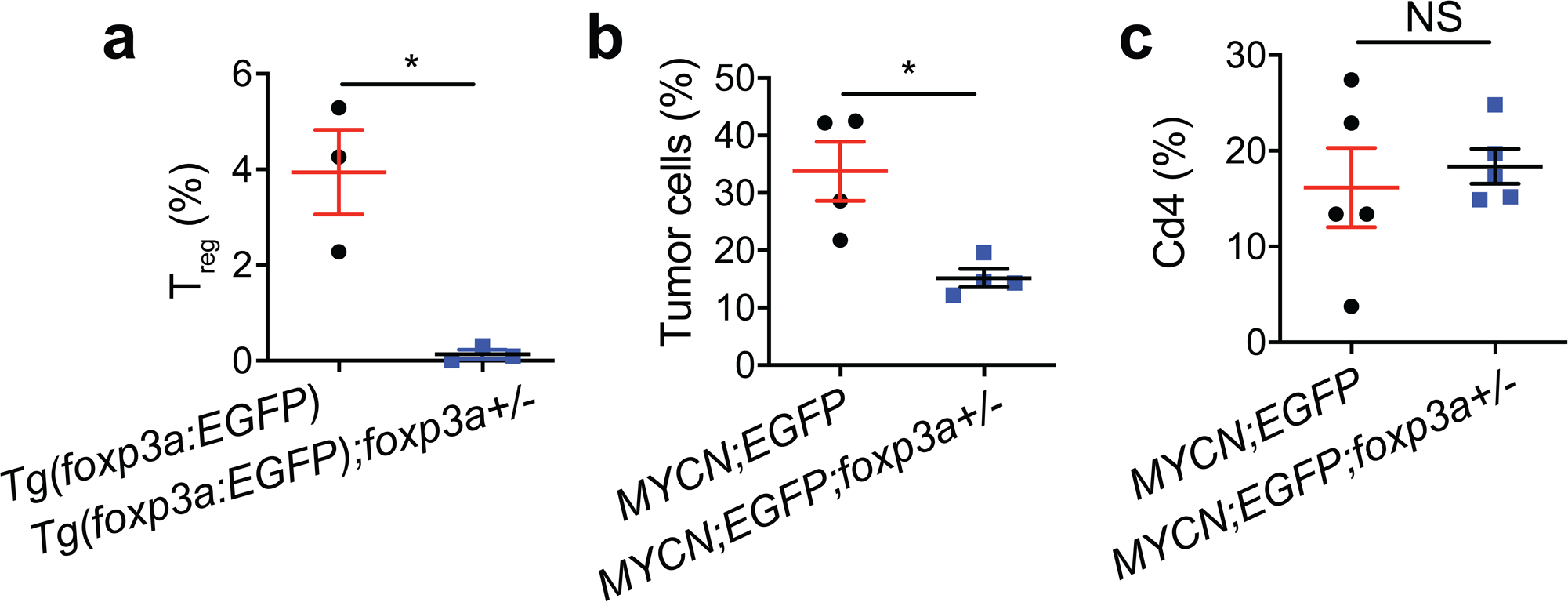
**a**, Flow cytometric analysis of the percentage of EGFP^+^ T_reg_ cells among the total lymphocytes inside the kidneys of *Tg(foxp3a:EGFP)* fish and their *Tg(foxp3a:EGFP);foxp3a+/-* siblings at six months of age (*n =* 3). **b**, Flow cytometric analysis of the percentage of EGFP^+^ tumor cells inside the TME of *Tg(foxp3a:EGFP)* fish and their *Tg(foxp3a:EGFP);foxp3a+/-* siblings at six months of age (*n =* 3). **c**, Flow cytometric analysis comparing Cd4^+^ cells from the kidney of *MYCN;foxp3a+/+ vs. MYCN;foxp3a+/-* sibling fish at six months of age (*n =* 3). Data are presented as mean ± SEM and analyzed for statistical significance by an unpaired two-tailed *t*-test. NS, not significant; * *P* < 0.05;

**Extended Data Fig. 2.**
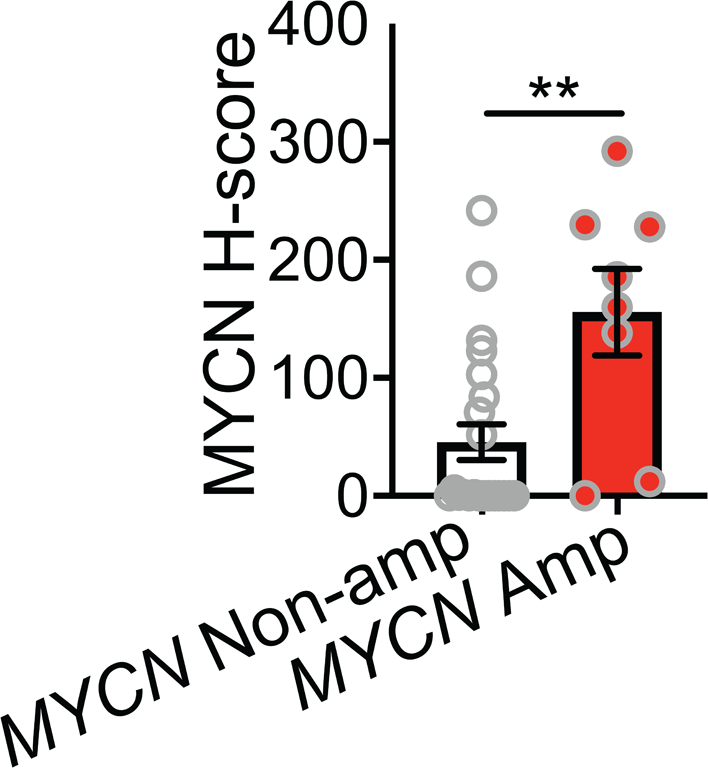
A significantly more MYCN protein levels in *MYCN*-amplified NB samples. Quantification of MYCN immunohistochemical staining as H-scores per tumor area in *MYCN* non-amplified (non-amp) and amplified (amp) NB patient samples (*n* = 22 for *MYCN* non-amp and *n* = 8 for amp tumor samples, respectively). *n* indicates the number of biological replicates. Data are presented as mean ± SEM and analyzed for statistical significance by an unpaired two-tailed *t*-test. ** *P* < 0.01.

**Extended Data Fig. 3.**
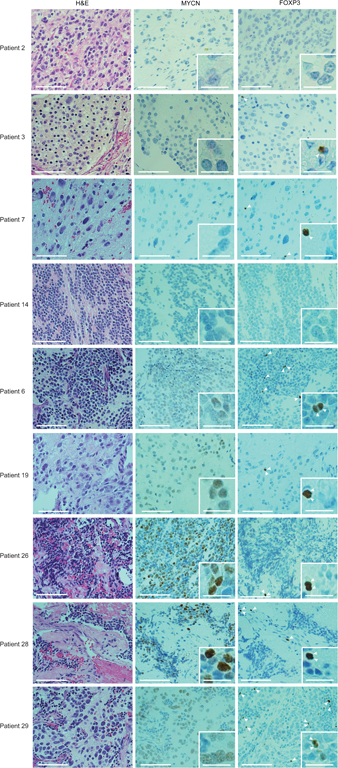
MYCN-high patient NB cells attract FOXP3^+^ T_reg_ cells. Immunohistochemical staining of MYCN and FOXP3 in primary NB patient samples. Scale bars = 200 and 50 (inserts) μm. Arrowheads point to FOXP3^+^ T cells.

